# Cytoplasmic HIF-2α correlates to proliferation and predicts worse outcome in sympathetic paraganglioma

**DOI:** 10.1101/2022.05.27.493680

**Authors:** Sinan Karakaya, Lisa Gunnesson, Erik Elias, Paula Martos Salvo, Mercedes Robledo, Ola Nilsson, Bo Wängberg, Frida Abel, Sven Påhlman, Andreas Muth, Sofie Mohlin

**Affiliations:** Division of Pediatrics, Department of Clinical Sciences, Lund University, Lund, Sweden; Lund Stem Cell Center, Lund University, Lund, Sweden; Lund University Cancer Center, Lund University, Lund, Sweden; Department of Surgery, Sahlgrenska University Hospital, Gothenburg, Sweden; Department of Surgery, Institute of Clinical Sciences, Sahlgrenska Academy, University of Gothenburg, Gothenburg, Sweden; Hereditary Endocrine Cancer Group, Spanish National Cancer Research Centre (CNIO), 28029 Madrid, Spain; Department of Laboratory Medicine, Institute of Biomedicine, Sahlgrenska Academy, University of Gothenburg, Gothenburg, Sweden; Department of Clinical Genetics and Genomics, Sahlgrenska University Hospital, Gothenburg, Sweden; Translational Cancer Research, Department of Laboratory Medicine, Lund University, Lund, Sweden

## Abstract

Pheochromocytomas (PCCs) and paragangliomas (PGLs) are rare neuroendocrine tumors. PGLs can further be divided into sympathetic (sPGLs) and head-and-neck (HN-PGLs). There are virtually no treatment options, and no cure, for metastatic PCCs and PGLs (PPGLs). Here, we composed a tissue microarray (TMA) consisting of 149 PPGLs, reflecting clinical features and presenting as a useful resource. Mutations in the pseudohypoxic marker *EPAS1*/HIF-2α correlates to an aggressive tumor phenotype. We show that HIF-2α unexpectedly localized to the cytoplasm in PPGLs. This subcompartmentalized protein expression differed between tumor subtypes, and strongly correlated to proliferation. Half of all sPGLs were metastatic at time of diagnosis. Cytoplasmic HIF-2α was strongly expressed in metastatic sPGLs and predicted poor outcome in this subgroup. We propose that cytoplasmic HIF-2α expression serves as a useful clinical marker to differentiate subtypes and predicting outcome, and hence can be used for improved targeted treatment in PPGLs.

## Introduction

Pheochromocytomas (PCCs) and paragangliomas (PGLs) are rare neuroendocrine tumors originating from schwann cell precursors in the adrenal medulla (PCCs)^1^, sympathetic ganglionic tissue in the abdomen or thorax (sympathetic PGL (sPGL)) or parasympathetic ganglionic tissue mainly in the head- and neck region (HN-PGLs).

Radical surgery is curative in most patients. Still, while most tumors display a non-metastatic phenotype, around 10% of pheochromocytomas and paragangliomas (PPGLs) develop metastases^2^. There are no reliable markers to conclusively demonstrate malignant potential of primary tumors. The 2017 WHO classification^3^ adopted the position that all PPGLs have malignant potential and instead made a distinction between non-metastatic and metastatic tumors. Since limited treatment options exist for patients with metastatic PPGL, better prognostic and predictive markers are needed for tailored treatment and follow-up.

PPGLs have the highest rate of heritability of all human solid tumors with up to 40% carrying germline mutations. The majority of familial PPGL tumors are caused by pathogenic variants in one of the four Succinate dehydrogenase subunit genes (*SDHx*). Sporadic PPGL harbor somatic driver mutations in some of the same genes as inherited predisposition syndromes (*e.g., RET, VHL, NF1, SDHB*), as well as in additional genes (*e.g., HRAS, EPAS1*)^4–8^.

Overall, there are over 20 susceptibility genes identified in PPGL^9^, and tumors can be divided into three broad groups; the pseudohypoxic cluster 1 with mutations in *VHL, SDHx, EGLN1/PHD2, FH* and *EPAS1* genes^10^, the kinase signaling cluster 2 with mutations in *RET, NF1, HRAS, MAX, TMEM127*, and the wnt pathway cluster 3 with *MAML3* fusions and mutations in *CSDE1*^8^. Mutations in the first group lead to increased stabilization of hypoxia inducible factor (HIF)-2α and a metabolic switch into a persistent pseudohypoxic state (*i.e*., hypoxic phenotype in oxygenated conditions), while mutations in the kinase signaling group of genes activates the PIK3/AKT/mTOR and RAS/RAF pathways^11,12^.

Oxygen sensing is one of the most important evolutionary traits of multicellular organisms (reviewed in Hammarlund et al.^13^). During physiological settings when oxygen is present, HIFs are targeted for degradation. There are however mechanisms by which HIFs escape this, *e.g*., loss-of-function mutations in the genes encoding VHL or PHD, generating pseudohypoxic phenotypes. While both HIF-1α and HIF-2α respond to oxygen fluctuations, it is primarily HIF-2α that has been implicated in tumor progression. For example, the majority of clear cell renal cell carcinomas are driven by mutations in *VHL*, leading to complete HIF-2-driven tumor formation^14^.

Ten years ago, the first mutation in *EPAS1* (encoding HIF-2α) in cancer was detected^15^. Two novel mutations were found in PGL patients, associated with increased protein half-life and HIF-2α activity^15^. Since then, a number of *EPAS1* mutations have been detected in PPGL^16–22^. Fliedner et al. further defined *EPAS1* mutated tumors as a subcluster of the pseuodohypoxic cluster 1 PGLs, separated from other hereditary pseudohypoxic PGLs driven by *VHL* or *SDHx* genes^23^. Recently, mutations in *EPAS1* have been detected also in esophageal squamous cell carcinoma and colorectal cancer^24,25^.

HIF-2α acts as a nuclear transcription factor. However, emerging evidence show that HIF-2α can localize to the cytoplasm in neuroblastoma^26,27^ and glioblastoma^28^. In similarity to PPGL, neuroblastoma arises along the sympathetic ganglia and in adrenal glands, and originates from stem cells of the neural crest. We recently showed that HIF-2α is expressed in the cytoplasm of migrating neural crest cells^29^, demonstrating the importance of understanding HIF-2α expression and biology in PPGL.

We provide here a comprehensive PPGL TMA with an in-depth characterization of the material. All sPGL specimens were positive for cytoplasmic HIF-2α and more than half of these patients presented with metastasis at time of diagnosis. Staining of Ki-67 as a proxy for proliferation is an important clinical parameter. We found that cytoplasmic HIF-2α strongly correlated with Ki-67, and high expression of cytoplasmic HIF-2α predicted poor outcome in sPGLs. Our results utilizing this TMA for biological analyses suggest that the cytoplasmic fraction of the HIF-2α protein separates metastatic from non-metastatic PPGL, confers aggressive features, and predicts outcome. We propose further studies to establish and implement cytoplasmic HIF-2α expression as a clinical marker for poor outcome in PPGL.

## Results

### Characterization of patient material

The TMA was constructed from donor blocks of fixated primary tumor tissue available from 149 out of a total of 175 patients diagnosed and operated for PPGL at Sahlgrenska University Hospital, Gothenburg, Sweden, between 1983 and 2015. The patient material (Table 1) is coherent with previously published population-based data^30^. The majority of patients were diagnosed with PCCs (n=122), while the rest were diagnosed with PGLs (n=27; 14 sPGLs and 13 HN-PGLs). At time of diagnosis, 93% (n=138) of the tumors were considered nonmetastatic while 7% (n=11) were diagnosed as metastatic. A substantially higher proportion of sPGLs were considered metastatic as compared to PCCs (50% vs. 3% respectively). None of the thirteen HN-PGLs were metastatic at time of diagnosis, in line with an overall low malignancy rate among these tumors^31^. In total, 44 patients (29%) were found to have somatic or germline mutations known to be associated with PPGL (Table 1), in coherence with previous data^32,33^.

**Table 1.**
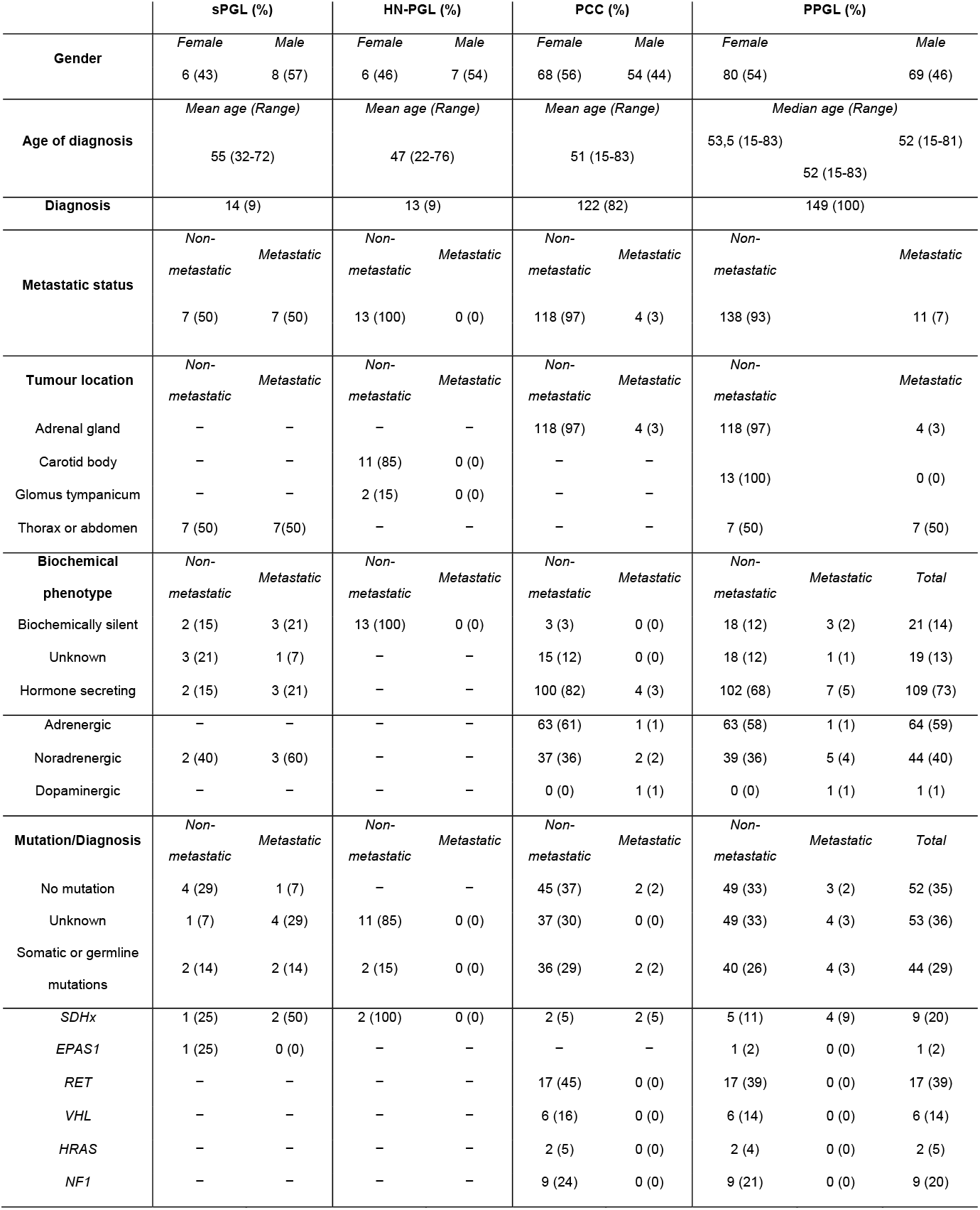
General characteristics of the patient cohort.

### PPGLs express clinically used diagnostic markers

We stained our TMA for diagnostic PPGL markers Chromogranin A (CgA), tyrosine hydroxylase (TH), vimentin (VIM), and synaptophysin (SYP) (Supplementary Fig. S1). All proteins were expressed as expected (Supplementary Fig. S1). None of these markers differentiated non-metastatic vs. metastatic PPGLs (Supplementary Table S2).

### sPGLs are more proliferative than PCCs

Generally, PPGLs are slow growing tumors with low proliferation rates, and Ki-67 positive tumor cells typically reside in hot spot regions. We stained and analyzed such hot spot regions from each TMA donor. The fraction of Ki-67 positive cells was significantly higher in sPGL as compared to PCCs (Fig. 1A-B and Table 2).

**Fig. 1.**
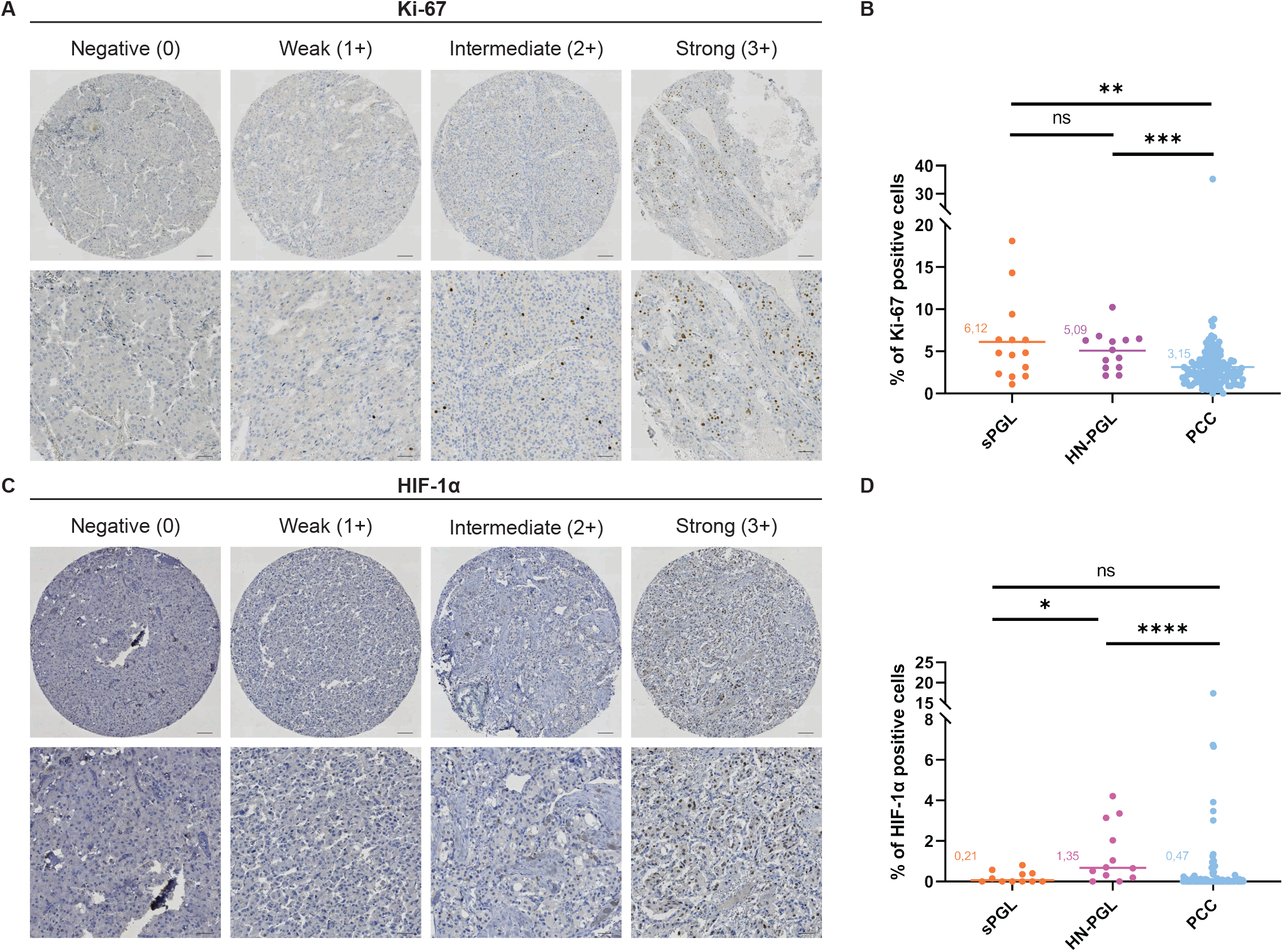
PPGLs express low levels of Ki-67 and HIF-1α. **A.** Representative images from tumor cores scored as negative (0), weak (1+), intermediate (2+) or strong (3+) for Ki67. **B.** Quantification of Ki67 positive cells (in %). **C.** Representative images from tumor cores scored as negative (0), weak (1+), intermediate (2+) or strong (3+) for HIF-1α. **D.** Quantification of HIF-1α positive cells (in %). Tumor cores were scored manually by three independent researchers as well as semi-automatically using the QuPath-0.3.0 software. Statistical significance was determined by the Mann-Whitney test. sPGL, sympathetic paraganglioma; HN-PGL, head- and neck paraganglioma; PCC, pheochromocytoma. Scale bar represents 100μm in core images and 50μm in zoomed images.

**Table 2.** Staining characteristics of the patient cohort.

| Marker | Diagnosis/Intensity | Negative (%) | Weak (%) | Intermediate (%) | Strong (%) | Positive (%) | Total # of Tumors (%) |
| --- | --- | --- | --- | --- | --- | --- | --- |
| **Ki-67** | *sPGL* | 0 (0) | 0 (0) | 4 (29) | 10 (71) | 14 (100) | 13 (100) |
|  | *HN-PGL* | 0 (0) | 0 (0) | 2 (16) | 11 (84) | 13 (100) | 13 (100) |
|  | *PCC* | 2 (2) | 13 (11) | 59 (48) | 48 (39) | 120 (98) | 122 (100) |
| **HIF-1α** | *sPGL* | 12 (100) | 0 (0) | 0 (0) | 0 (0) | 0 (0) | 12 (100) |
|  | *HN-PGL* | 8 (62) | 5 (38) | 0 (0) | 0 (0) | 5 (38) | 13 (100) |
|  | *PCC* | 105 (91) | 7 (6) | 1 (1) | 3 (2) | 11 (9) | 116 (100) |
| **HIF-2α-N** | *sPGL* | 8 (62) | 4 (31) | 1 (7) | 0 (0) | 5 (38) | 13 (100) |
|  | *HN-PGL* | 1 (8) | 7 (58) | 1 (8) | 3 (26) | 11 (92) | 12 (100) |
|  | *PCC* | 97 (82) | 19 (16) | 2 (2) | 0 (0) | 21 (18) | 118 (100) |
| **HIF-2α-C** | *sPGL* | 0 (0) | 4 (31) | 7 (54) | 2 (15) | 13 (100) | 13 (100) |
|  | *HN-PGL* | 0 (0) | 1 (8) | 5 (42) | 6 (50) | 12 (100) | 12 (100) |
|  | *PCC* | 15 (13) | 64 (54) | 33 (28) | 6 (5) | 103 (87) | 118 (100) |

### PPGLs express low levels of HIF-1α

Since a large fraction of PPGLs presents with mutations affecting oxygen sensing, we stained for HIF-1α and HIF-2α. While 38% of HN-PGLs were positive, only 9% of PCCs and 0% of sPGLs expressed HIF-1α (Fig. 1C-D and Table 2). There was no difference in HIF-1α staining between non-metastatic and metastatic tumors (Supplementary Table S2).

### HIF-2α expression is mainly cytoplasmic in all paragangliomas

HIF-2α positive tumor cores showed a high degree of cytoplasmic staining (Fig. 2A and Table 2), in concordance with findings in glioblastoma^28^, neuroblastoma^27^, and neural crest development^29^. We therefore decided to divide our analyses into nuclear and cytoplasmic HIF-2α. Evaluation of staining intensity of HIF-2α revealed that the majority of tumors scored negative for nuclear HIF-2α, while highly intense cytoplasmic HIF-2α staining was observed in 69% of sPGLs, 92% of HN-PGLs and 33% of PCCs (Fig. 2B-E and Table 2). To address HIF-2 activity in these tumors, we stained for HIF-2 specific target gene DEC1/BHLBE40, and found that the majority of tumors from all subtypes expressed this protein (Supplementary Table S2).

**Fig. 2.**
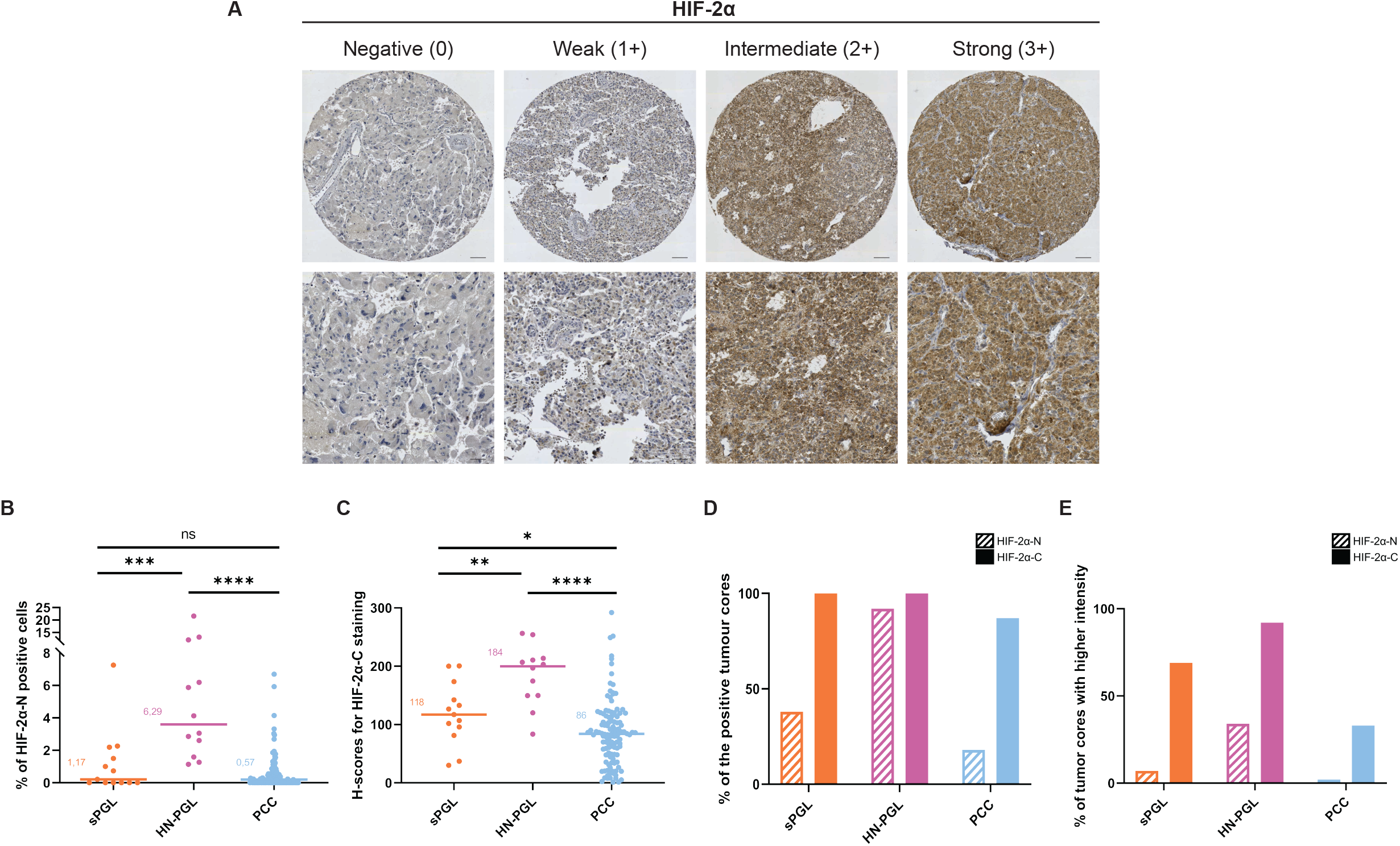
PPGLs express high levels of cytoplasmic HIF-2α. **A.** Representative images from tumor cores scored as negative (0), weak (1+), intermediate (2+) or strong (3+) for HIF-2α. **B.** Quantification of positive cells for nuclear HIF-2α (HIF-2α-N) (in %). **C.** H-score for cytoplasmic HIF-2α (HIF-2α-C). **D.** Quantification of fraction (in %) of positive tumor cores for nuclear and cytoplasmic HIF-2α. **E.** Quantification of fraction (in %) of tumor cores staining with high intensity (scored 2-3) for nuclear and cytoplasmic HIF-2α. Tumor cores were scored manually by three independent researchers as well as semi-automatically using the QuPath-0.3.0 software. Statistical significance was determined by the Mann-Whitney test. sPGL, sympathetic paraganglioma; HN-PGL, head- and neck paraganglioma; PCC, pheochromocytoma. Scale bar represents 100μm in core images and 50μm in zoomed images.

### Metastatic sPGLs express high levels of cytoplasmic HIF-2α

Expression of nuclear HIF-2α was unchanged in non-metastatic vs. metastatic tumors in all three PPGL entities (Fig. 3A). In coherence with cytoplasmic HIF-2α staining at low intensity in PCCs overall, there was no difference in expression in non-metastatic as compared to metastatic tumors (Fig. 3B and Supplementary Table S1). Notably, while 43% of nonmetastatic sPGLs strongly expressed cytoplasmic HIF-2α, this number increased to 83% in metastatic tumors (Fig. 3B and Supplementary Table S2).

**Fig. 3.**
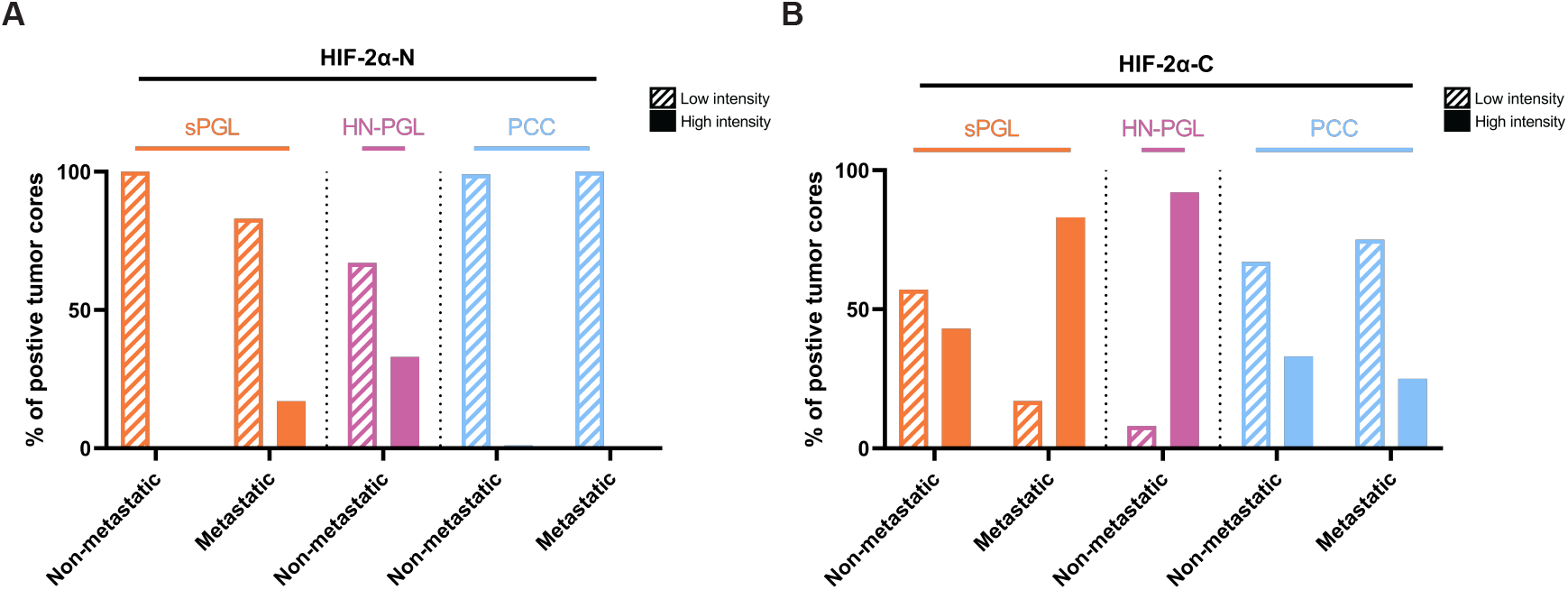
Cytoplasmic HIF-2α is highly expressed in metastatic sPGLs. **A-B.** Expression of nuclear (HIF-2α-N) (A) and cytoplasmic (HIF-2α-C) HIF-2α as divided by low (0-1) and high (2-3) staining intensity in sPGL, HN-PGL and PCC tumor subgroups. Note that there are no metastatic HN-PGLs in this cohort. sPGL, sympathetic paraganglioma; HN-PGL, head- and neck paraganglioma; PCC, pheochromocytoma.

### Cytoplasmic HIF-2α correlates to proliferation proxy Ki-67

To determine the putative clinical impact of cytoplasmic HIF-2α, we investigated possible relationships with PPGL diagnostic markers. When investigating PPGLs as one entity, we found that cytoplasmic HIF-2α positively correlated with SYP, DEC1/BHLHE40 and Ki-67, while there was an inverse correlation with CgA expression (Fig. 4A-B). Correlation between HIF-2α and Ki-67 was strongest in sPGLs (Fig. 4C).

**Fig. 4.**
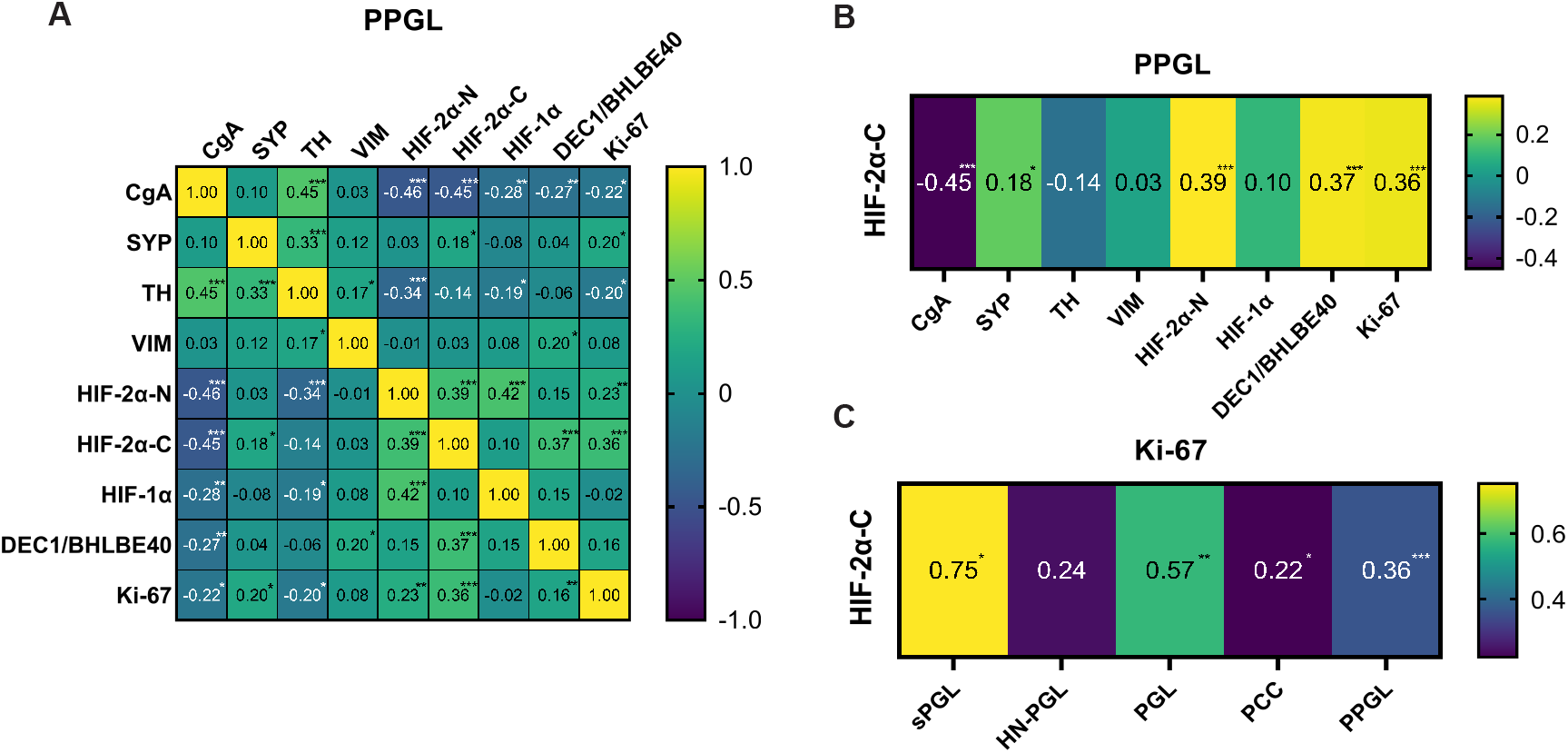
Cytoplasmic HIF-2α correlates to proliferation in sPGLs. **A.** Compilation of correlations between all markers stained for in this cohort. **B.** Compilation of correlations only between cytoplasmic HIF-2α and the other markers for visualization. **C.** Correlations between cytoplasmic HIF-2α and Ki-67 as divided into the different subgroups of PPGL. Correlation coefficient was calculated using Spearman’s correlation test in GraphPad Prism 9 and statistical significance (two-tailed P value for each correlation coefficient) was computed automatically by the software. sPGL, sympathetic paraganglioma; HN-PGL, head- and neck paraganglioma; PGL, paraganglioma; PCC, pheochromocytoma; PPGL, pheochromocytoma and paraganglioma.

### EPAS1 mutated tumors express cytoplasmic HIF-2α

We analyzed expression of HIF-2α in five tumors with *EPAS1* mutations from two different cohorts (one tumor from TMA; four tumors from the CNIO study cohort^34^). All of these tumors (1 PCC and 4 sPGLs) expressed cytoplasmic HIF-2α (Supplementary Fig. S2 and Supplementary Table S3).

### EPAS1 expression is higher in sPGLs

We have previously performed whole exome sequencing on a subset of PPGLs included in our TMA^35^. Here we utilized this material to analyze *HIF1A* and *EPAS1* expression. Expression of *HIF1A* mRNA was slightly higher in the pseudohypoxic cluster, but half of the tumors in this group still displayed low levels (Fig. 5A). We could not detect any correlations of *HIF1A* mRNA and mutation status, metastasis or diagnosis (Fig. 5A). Analysis of *EPAS1* expression showed substantially higher expression levels in tumors belonging to the pseudohypoxic cluster (Fig. 5A-B). We did not find a correlation between mRNA and cytoplasmic HIF-2α expression as most tumors in the kinase signaling cluster express the protein, but low mRNA levels (Fig. 5A, C).

**Fig. 5.**
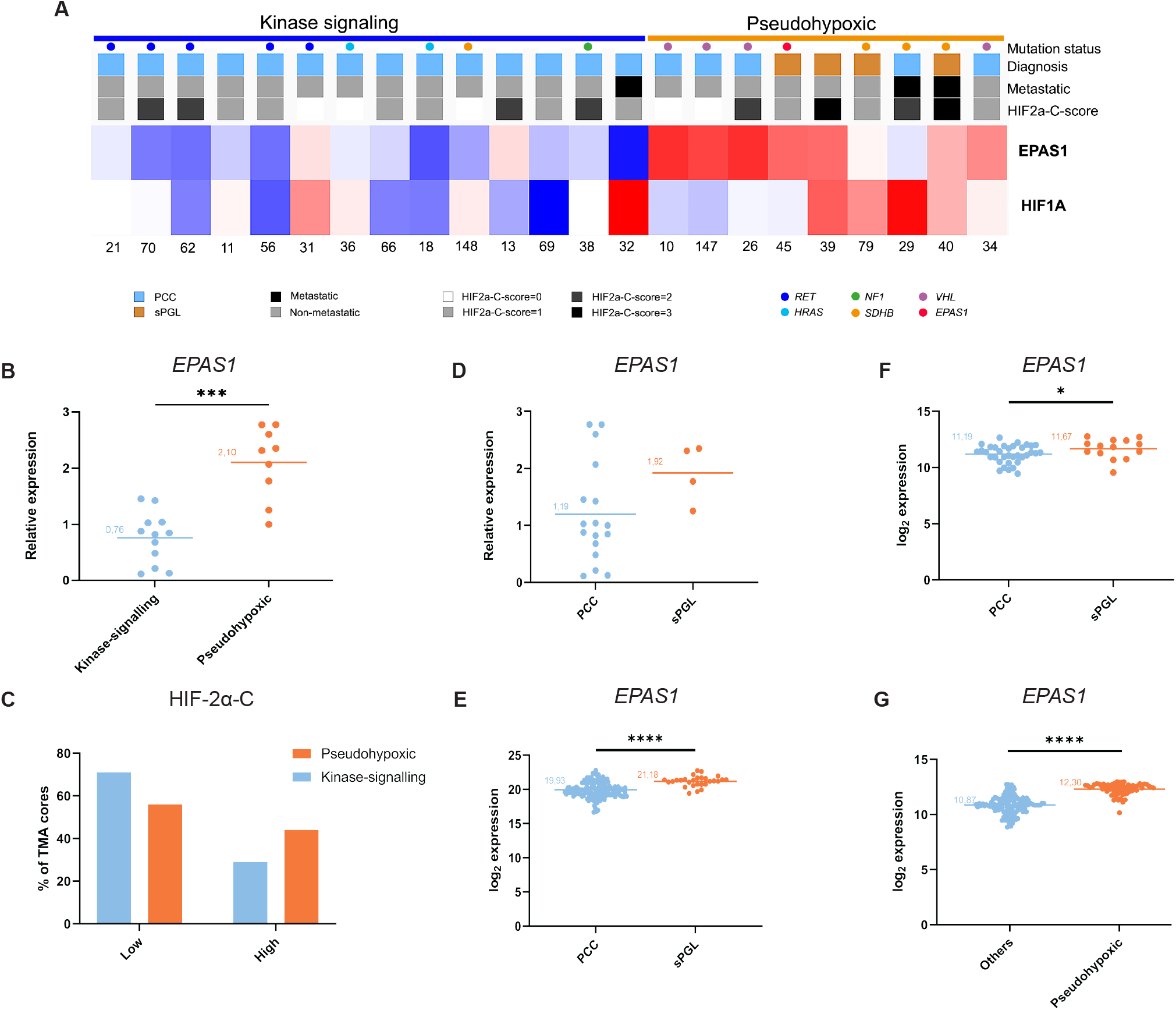
EPAS1 mRNA expression is higher in sPGL as compared to PCC. **A.** *HIF1A* and *EPAS1* mRNA expression in a subset of tumors from our TMA cohort as analyzed by whole exome sequencing. **B.** Expression of *EPAS1* mRNA in the pseudohypoxic and kinase-signaling groups from (A). **C.** Quantification of fraction of TMA cores scored as low or high for cytoplasmic HIF-2α in the pseudohypoxic and kinase-signaling groups. **D-F.** Expression of *EPAS1* mRNA as determined by RNA sequencing in sPGLs and PCCs in three different cohorts; TMA (n= 21, **D**), TCGA (n=177, **E**) and Korpershoek (n=49, **F**). **G.** Expression of *EPAS1* mRNA as determined by RNA sequencing in pseudohypoxic as compared to all other subgroups in the Favier cohort (n=188). Statistical significance was determined using Mann-Whitney test. PCC, pheochromocytoma; sPGL, sympathetic paraganglioma.

We further investigated mRNA expression in correlation to tumor subtype and found that *EPAS1* levels were higher in sPGLs as compared to PCCs (Fig. 5D), in line with protein data (Fig. 2). We confirmed these findings in two independent cohorts (TCGA, n=177 and Korpershoek, n=49; Fig. 5E-F). While we could not assess *EPAS1* expression between PPGL subtypes in the Favier data set (n=188) due to lack of such data, we detected higher mRNA levels in pseudohypoxic tumors as compared to all other groups (Fig. 5G).

### sPGL patients die of metastatic disease

To investigate the survival status in our cohort, we used clinical follow-up data where 25 patients had died at an end point of 10 years (Supplementary Table S4). Dividing the cohort according to diagnosis, 15% of PCC, and as many as 50% of sPGL patients were deceased. Out of these 25 patients in total, 9 had died from cancer (5 from metastatic PPGL and 4 from other cancers), while the remaining 16 patients died of other causes (heart-related, infections, multi-organ failure, stroke, or other or unknown reasons). The survival probability did not differ between PCC and PGL patients when taking all causes of death into consideration (Supplementary Fig. S3A). However, when further dividing into PGL subtypes, sPGLs displayed a significantly worse prognosis than PCCs and HN-PGLs (Fig. 6A). Since it is difficult to determine whether the cause of death from other reasons than metastatic PPGLs were due to their primary tumor, we stratified survival probability only in patients that died from metastatic disease. PGLs present with worse survival than PCC (Supplementary Fig. S3B). Although few patients in this cohort died from their disease, when further stratifying PGL patients, there is a clear separation where sPGL patients have worse prognosis (survival rate only 68% for sPGL, while 100% for HN-PGL and 98% for PCC) (Fig. 6B).

**Fig. 6.**
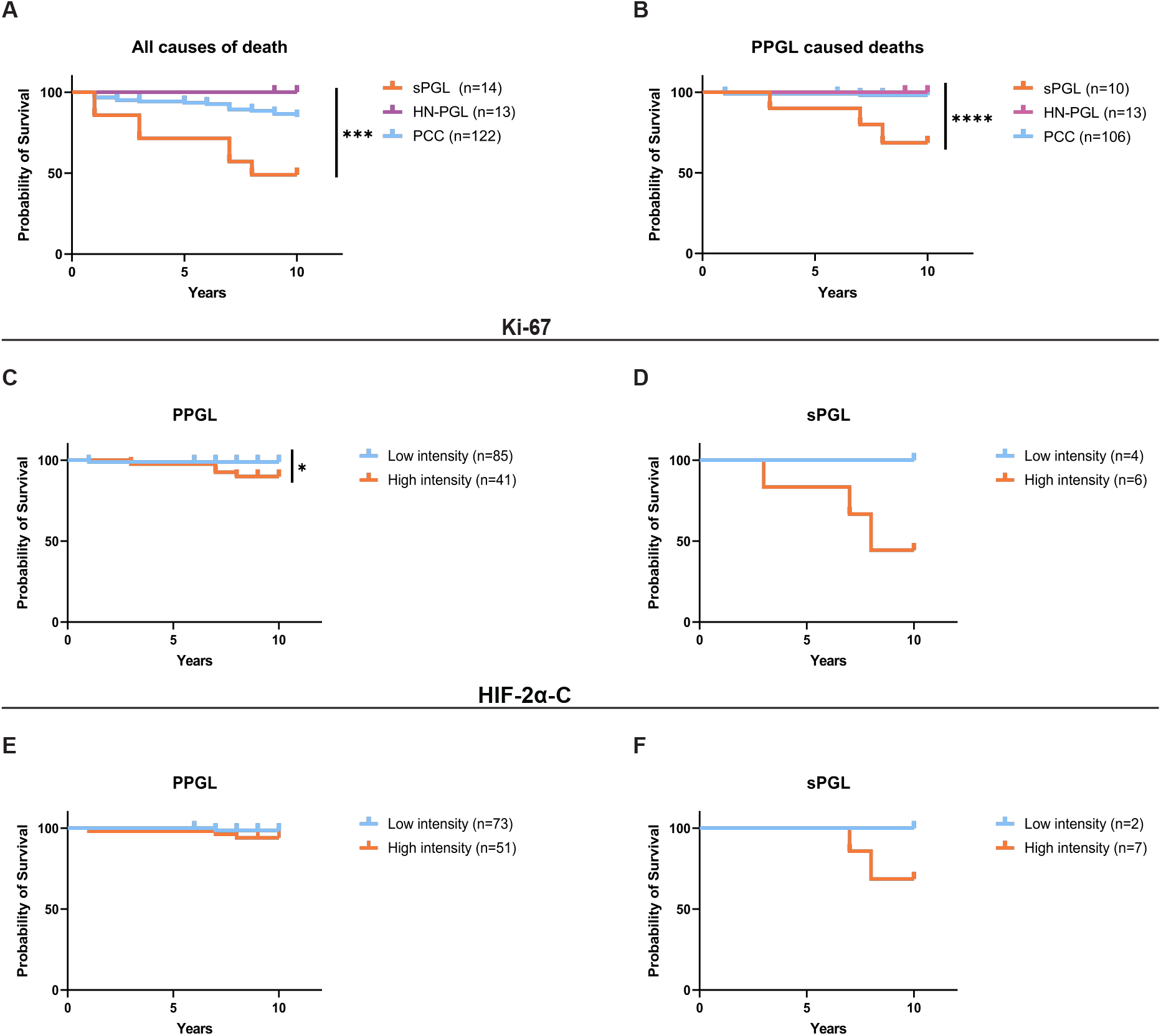
Intense staining of cytoplasmic HIF-2α predicts poor outcome in sPGLs. **A.** Survival probability (10-year follow-up) of PPGL patients when including all causes of death. **B.** Survival probability (10-year follow-up) of patients when only including patients dead from PPGL disease comparing PCC and PGL subgroups. **C-D**. Survival probability stratified by low (0-1) and high (2-3) staining intensity of Ki-67 in the entire PPGL cohort (**C**) and sPGLs only (**D**). **E-F**. Survival probability stratified by low (0-1) and high (2-3) staining intensity of cytoplasmic HIF-2α in the entire PPGL cohort (**E**) and sPGLs only (**F**). Statistical significance was calculated using the Log-rank (Mantel-Cox) test. PCC, pheochromocytoma; sPGL, sympathetic paraganglioma; HN-PGL, head- and neck paraganglioma. PGL, paraganglioma; PPGL, pheochromocytoma and paraganglioma.

### High expression of Ki-67 and cytoplasmic HIF-2α predicts worse survival in sPGL patients

There was no difference in survival between patients with low or high Ki-67 expression in the PCC entity (Supplementary Fig. S3C). However, high intensity of Ki-67 positivity predicts worse outcome when analyzing the entire cohort (Fig. 6C), explained by a separation in survival in PGL patients (Supplementary Fig. S3D), and an even more pronounced effect when sPGL patients were analyzed separately (Fig. 6D). While there was no correlation between cytoplasmic HIF-2α and outcome in PPGL or PCC entities (Fig. 6E and Supplementary Fig. S3E), high intensity of cytoplasmic HIF-2α predicts poor outcome in PGL patients (Supplementary Fig. S3F), with a more pronounced effect in sPGL patients specifically (Fig. 6F).

## Discussion

There is a lack of accessible material for biological analyses of PPGL. We present a TMA of cores from 149 patients with PCCs, HN-PGLs and sPGLs, reflecting patient distribution of all clinical parameters. We demonstrate that this material is valuable by extensive characterization of protein markers used in diagnostics in connection to primary tumor location, non-metastatic vs. metastatic disease, mutation status, biochemical phenotype, and importantly 10-year follow-up data. In addition, we add RNA expression data from a selection of these patients.

Half of sPGL patients in our cohort present with metastatic disease, and in coherence, patients from this tumor entity have a worse prognosis and lower survival rate as compared to PCC and HN-PGL. This strengthens the benefit of the WHO classification of these three subgroups for diagnostics, when assessing treatment regimens and patient follow-up schemes. It has recently been further demonstrated that metastatic HN-PGLs have a longer disease-specific survival than metastatic PPGLs, and that extra-adrenal PPGLs (i.e., PGL as compared to PCC) have worse prognosis^36^, in line with our data.

PPGLs within the pseudohypoxic group present with a more aggressive behavior and worse prognosis. Mutations in the *EPAS1* gene confer such a pseudohypoxic profile and represent a subgroup of its own within the pseudohypoxic group of tumors^23^. We have, in previous studies on neural crest cells during embryogenesis as well as in neuroblastoma, observed that the HIF-2α protein non-canonically localizes to the cytoplasm^27,29^. There was a clear distinction between the percentage of tumors positive for cytoplasmic vs. nuclear protein in PPGLs, and considering their proposed shared ancestor cells, the neural crest, with neuroblastoma (shown in e.g., composite tumors^37^), these findings are of importance to further understand PPGL initiation and subtype distinction.

The relatively high expression of HIF-1α and nuclear and cytoplasmic HIF-2α in HN-PGLs is explained by the fact that these tumors are located in the oxygen sensing carotid body. In contrast, sPGLs express no or very low levels of HIF-1α and nuclear HIF-2α, while all of them express substantial levels of cytoplasmic HIF-2α. This suggests that the presence of cytoplasmic HIF-2α in sPGLs is not due to a hypoxic environment in these tumors, but rather present with a bona fide pseudohypoxic phenotype mediated via non-canonical mechanisms for regulation of HIF-2α and downstream activity.

PPGLs are to a large extent slow growing, and the assessment of Ki-67 expression as a proxy for proliferation in clinical assessment of these tumors is of significant value for prognostics. Sympathetic PGLs display substantially higher expression of Ki-67 and in coherence with this more proliferative phenotype predicts poor outcome for patients that die from their primary tumor. Cytoplasmic HIF-2α correlates positively to Ki-67, and is highly and intensely expressed in the majority of metastatic sPGLs cases. Albeit few patients die from their primary disease, while all sPGL patients that present with low tumor intensity of cytoplasmic HIF-2α survive, the fraction of patients with high intensity-expressing tumors presents with a survival probability of less than 50%.

The mechanism-of-action for cytoplasmic HIF-2α is yet unknown, and ongoing functional studies aim to assess its direct role in PPGL subtype initiation and progression. Our data show that *EPAS1* mRNA expression is substantially increased in the pseudohypoxic-as compared to the kinase PPGL subgroup. However, both groups express high cytoplasmic HIF-2α, suggesting that HIF-2α is regulated at a (post-)translational-rather than transcriptional level. These results are in line with previously published data where expression of classical HIF-2 transcriptional target genes did not differ between the PPGL subclusters, suggesting non-canonical functions of HIF-2α^38^.

To assess the role of HIF-2α in PPGL progression, Bechmann and colleagues knocked down or overexpressed HIF-2α in cell line models, and showed that HIF-2α induced a prometastatic phenotype^39^. The HIF-2α-driven effects on the PPGL cells were due to protein function per se and not hypoxia-induced mechanisms^39^. These data on overall HIF-2α protein expression in PPGL as one tumor entity, in combination with our results on the dependence of cytoplasmic HIF-2α in the different tumor subtypes, strongly suggest an important tumor-driving role of HIF-2α in these tumors.

sPGLs and NBs arise along the sympathetic chain ganglia and share a common ancestor cell in the trunk neural crest. It is therefore of particular interest that the HIF-2α protein is expressed in oxygenated cells, is mainly cytoplasmic, and that this fraction of cells predicts poor outcome in both of these tumor entities. In addition, the high expression of cytoplasmic HIF-2α in metastatic sPGLs is coherent with the correlation between intensely HIF-2α positive tumor cells and distal metastasis in NB. Since sPGL patients present with significantly worse survival than PCC and HN-PGL patients, routinely assessing HIF-2α expression in sPGL could represent a new diagnostic biomarker and aid in treatment decisions and patient follow-up.

## Material and Methods

### Clinical Material

A total of 149 patients diagnosed with PPGL at Sahlgrenska University Hospital, Gothenburg, Sweden, between 1984 and 2015 were included in the TMA. Patient information concerning age, sex, recurrent or metastatic disease as well as hormone profile of the tumors and causes of death when applicable was retrieved from in-patient charts and a pre-existing clinical database. Clinical pathology reports were searched for any further tumor specific data. Characterization of the material is summarized in Table 1.

### Tissue microarray and immunohistochemical staining

Paraffin blocks of paragangliomas or pheochromocytomas were retrieved from the pathology archives at Sahlgrenska University Hospital, Gothenburg. From representative tumor areas selected by a board-certified pathologist (O.N.), tissue core biopsies were punched out and mounted in recipient blocks. A total of 175 donor tumor cores with 149 unique primary tumors were included in the TMA. Sufficient core quality was assessed by eosin staining. Sections (4μm) were stained using AutostainerPlus (Dako). Detailed information on antibodies and scoring procedure can be found in *Online Methods*.

### Genetics

Germline mutational data of *SDHA*, *SDHB*, *SDHC*, *SDHD*, *RET*, *VHL*, *NF1* were collected from the two clinical genetics laboratories and oncogenetic clinics at the University hospitals in Gothenburg and Lund. Genetic screening of the proband/affected family member was performed using Sanger DNA sequencing or massive parallel sequencing panels, and for patients diagnosed from 2014 and onwards the analyses were combined with multiplex ligation-dependent probe amplification (MLPA, P081-D1 & P082-C2 (NF1), P226-D1 (SDHx), and P016-C2 (VHL), MRC-Holland, Amsterdam, The Netherlands) for the detection of large deletions or duplications.

Whole exome sequencing (WES) data from 23 PPGL cases was extracted from previous published papers^9,35^. WES was run as either paired tumor tissue and normal tissue/blood (10 cases) or as single tumor tissue (13 cases), and identified additional germline and somatic variants in PPGL susceptibility genes (i.e., *NF1, VHL, RET, EPAS1* and *HRAS*). All tumor samples with no apparent causative point-mutation were further analyzed for exon/gene deletions or duplications by MLPA as described above. Only variants predicted to be pathogenic or likely pathogenic by American College of Medical Genetics and Genomics^40^ were included in the final mutation list (Table 1)^40^.

### mRNA expression analyses

For 23 PPGL samples analyzed in the current study, mRNA expression data from fresh tumor tissue run by 44K Agilent Cy3/Cy5 2-color microarrays (Agilent) were available. Tumor samples were divided into expression groups based on previous unsupervised hierarchical clustering using a 153 discriminative gene set^35^.

Normalized and log transformed expression data from different PPGL data sets (TCGA, Loriot et al. (E-MTAB-733), Evenepoel et al. (GSE67066)) were downloaded from ‘R2: Genomics Analysis and Visualization Platform’.

### Statistics

GraphPad Prism 9 software was used for correlation, statistical and survival analyses. Mann-Whitney test was used to determine statistical significance and p-value was employed as; p<0.05 (*), p<0.01(**), p<0.001(***) and p<0.0001(****) if not otherwise specified. Spearman’s correlation test was used to calculate correlation coefficient and statistical significance (two-tailed P value for each correlation coefficient). Kaplan-Meier survival analysis and the Log-rank (Mantel-Cox) test were used to prepare and compare survival curves, respectively.

## Online Methods

### Immunohistochemistry

The TMA was stained for chromogranin A (CgA, MAB319, Merck Millipore 1:4000), synaptophysin (SYP, M0776, Agilent, 1:200+Linker), tyrosine hydroxylase (TH, ab75875, Abcam, 1:300), vimentin (ab45939, Abcam, 1:200), HIF-1α (610959, BD Biosciences, 1:50 + Linker), HIF-2α (BL-95-1A2, Nordic Biosite, 1:200+Linker), DEC1/BHLHE40 (nb100-1800, Novus Biologicals, 1:100), and Ki-67 (IR62661, Agilent, Ready-to-use + Linker). The slides were digitalized with Axio Scan Z.1 at 20x magnification.

### TMA scoring

Immunohistochemical (IHC) staining was viewed and examined blindly by two independent persons using the PathXL software. A second independent assessment was conducted by an independent person using QuPath-0.3.0 software^41^. The software’s TMA dearrayer was applied to each slide to detect individual tumor cores and manually corrected when necessary. For CgA, SYP, TH, VIM and HIF-2α-C a scoring intensity of negative (0), weak (1+), median (2+) and strong (3+) tumor cell specific expression was manually determined (representative images in Supplementary Fig. S1). Additionally, H-Score between 0-300 was calculated for HIF-2α-C staining for each tumor core by using QuPath-0.3.0 software. For this, cell detection command was used to identify each cell based on nuclear hematoxylin staining in every tumor core, then, intensity thresholds were set to distinguish cells with negative, weak, intermediate, and strong cytoplasmic HIF-2α based on DAB staining. The cores having H-Score between 0-15 scored as negative (0), 16-99 scored as weak (1+), 100-199 scored as intermediate (2+) and 200-300 scored as strong (3+).Four *EPAS1* mutated tumors (1 PCC, 3 sPGL, all non-metastatic) from the CNIO cohort^34^ were stained for HIF-2α using the same protocol as above. Scoring was performed by an independent person using the QuPath-0.3.0 software as described above. For HIF-1α, DEC1/BHLHE40 and nuclear HIF-2α staining, tumor cell specific expression scoring was determined semi-automatically by using QuPath-0.3.0 software. Each TMA core and cells were detected as above and then tumor cells with nuclear DAB staining were manually counted. The tumor cores with positive cancer cell percentages between 0-1% scored as negative (0), 1-5% scored as weak (1+), 5-10% scored as intermediate (2+) and >10% scored as strong (3+).

For the Ki-67 staining, the number of positive and negative cells in the hotspot regions detected by a pathologist were manually counted and percentage of positive cells were calculated for each tumor core. The tumor cores with positive cancer cell percentages between 0% scored as negative (0), 0-1% scored as weak (1+), 1-3% scored as intermediate (2+) and >3% scored as strong (3+)^42^. Cells of necrosis, healthy tissue, scanning and staining artifacts and folded areas were manually removed from the calculation.

### Specificity control of HIF-2a antibody for IHC

Rabbit anti-HIF-2α antibody from Nordic Biosite (BL-95-1A2, 1:200+Linker) was used to stain the TMA using the same protocol as above. Specificity was confirmed by staining hypoxic (1% oxygen) and normoxic (21% oxygen) SK-N-BE(2)c neuroblastoma, HEP3B hepatocellular carcinoma, and VHL negative renal cell carcinoma 786-O and RCC-4 cells.

## Funding

This work was supported by The Mrs Berta von Kamprad’s Foundation; the Swedish Cancer Society; the Swedish Childhood Cancer Fund; the Lions Cancer Fund West; and institutional research funds at the Department of Surgery, Sahlgrenska University Hospital.

## Notes

## Acknowledgements

The authors would like to thank Siv Beckman, Christina Möller, Gülay Altiparmak, and Yvonne Arvidsson for excellent technical assistance.

## Author contributions

Conceptualization: SK, LG, EE, BW, FA, SP, AM, SM; Data curation: SK, LG, FA; Formal analysis: SK, LG; Investigation: SK, LG, FA, ON, SP; Methodology: SK, LG, FA, SP, AM, SM; Resources: RM, ON, BW, FA, AM; Validation: SK, LG, FA, SP; Visualization: SK, FA; Writing—original draft: SM; Writing—review & editing: SK, LG, EE, PMS, RM, ON, BW, FA, SP, AM, SM; Supervision: AM, SM; Project administration: AM, SM; Funding acquisition: BW, AM, SM.

## Author disclosures

The authors have no conflicts of interest to report.

## Data Availability

The data underlying this article will be shared on reasonable request to the corresponding author.

## Supplementary material

**Fig. S1.** *PPGLs express clinically relevant markers*. Representative images from tumor cores scored as negative (0), weak (1+), intermediate (2+) or strong (3+) for PPGL diagnostic markers CgA, TH, SYP, Vimentin, as well as HIF-2 downstream marker DEC1/BHLHE40. Scale bar represents 100μm.

**Fig. S2.** *EPAS1 mutated tumors express cytoplasmic HIF-2α*. Images from tumor cores from four individual patients with mutations in the *EPAS1* gene from the CNIO cohort^34^ stained for HIF-2α. Scale bar represents 50μm.

**Fig. S3.** *Ki-67 or cytoplasmic HIF-2α expression does not stratify patient outcome in PCCs*. **A-B.** Survival probability (10-year follow-up) of PPGL patients divided into PGL and PCC subgroups when including all causes of death **(A)** and only PPGL caused deaths **(B)**. **C-D.** Survival probability (10-year follow-up) of patients when only including patients dead from PPGL disease in PCC (**C**) and PGL (**D**), comparing low (0-1) and high (2-3) staining intensity of Ki-67. **E-F**. Survival probability (10-year follow-up) of patients when only including patients dead from PPGL disease in PCC (**E**) and PGL (**F**) comparing low (0-1) and high (2-3) staining intensity of cytoplasmic HIF-2α.

**Table S1.** CgA, TH, SYP, VIM, DEC1/BHLBE40 staining characteristics of the cohort.

| Marker | Diagnosis/Intensity | Negative (%) | Weak (%) | Intermediate (%) | Strong (%) | Positive (%) | Total # of Tumors (%) |
| --- | --- | --- | --- | --- | --- | --- | --- |
| **CgA** | *sPGL* | 3 (23) | 4 (31) | 4 (31) | 2 (15) | 10 (77) | 13 (100) |
|  | *HN-PGL* | 6 (46) | 5 (38) | 2 (16) | 0 (0) | 7 (54) | 13 (100) |
|  | *PCC* | 1 (1) | 11 (9) | 24 (20) | 83 (70) | 118 (99) | 119 (100) |
| **TH** | *sPGL* | 6 (46) | 0 (0) | 2 (16) | 5 (38) | 7 (54) | 13 (100) |
|  | *HN-PGL* | 13 (100) | 0 (0) | 0 (0) | 0 (0) | 0 (0) | 13 (100) |
|  | *PCC* | 3 (3) | 9 (8) | 13 (11) | 94 (78) | 116 (97) | 119 (100) |
| **SYP** | *sPGL* | 0 (0) | 3 (23) | 5 (38) | 5 (38) | 13 (100) | 13 (100) |
|  | *HN-PGL* | 0 (0) | 0 (0) | 1 (8) | 12 (92) | 13 (100) | 13 (100) |
|  | *PCC* | 1 (1) | 6 (5) | 14 (12) | 98 (82) | 119 (100) | 119 (100) |
| **VIM** | *sPGL* | 3 (23) | 1 (8) | 3 (23) | 6 (46) | 10 (77) | 13 (100) |
|  | *HN-PGL* | 0 (0) | 0 (0) | 4 (31) | 9 (69) | 13 (100) | 13 (100) |
|  | *PCC* | 13 (11) | 14 (12) | 23 (19) | 69 (58) | 105 (89) | 119 (100) |
| **DEC1/BHLHE40** | *sPGL* | 3 (23) | 2 (15) | 0 (0) | 8 (62) | 10 (77) | 13 (100) |
|  | *HN-PGL* | 0 (0) | 1 (8) | 0 (0) | 12 (92) | 13 (100) | 13 (100) |
|  | *PCC* | 29 (24) | 30 (25) | 13 (11) | 47 (40) | 90 (76) | 119 (100) |

**Table S2.** HIF-2α staining in non-metastatic vs metastatic disease. Number and fraction (%) of cores staining with low (0-1) or high (2-3) intensity for nuclear (HIF-2α-N) or cytoplasmic (HIF-2α-C) HIF-2a in non-metastatic vs. metastatic tumors. PCC, pheochromocytoma; sPGL, sympathetic paraganglioma; HN-PGL, head- and neck paraganglioma.

| Marker | Intensity | Low intensity cores |  | High intensity cores |  | Total # of Tumors |  |
| --- | --- | --- | --- | --- | --- | --- | --- |
|  | Diagnosis | Non-metastatic (%) | Metastatic (%) | Non-metastatic (%) | Metastatic (%) | Non-metastatic (%) | Metastatic (%) |
| CgA | sPGL | 4 (66) | 3 (43) | 2 (34) | 4 (57) | 6 (100) | 7 (100) |
|  | HN-PGL | 11 (85) | – | 2 (15) | – | 13 (100) | – |
|  | PCC | 10 (9) | 2 (50) | 105 (91) | 2 (50) | 115 (100) | 4 (100) |
| TH | sPGL | 3 (50) | 3 (43) | 3 (50) | 4 (57) | 6 (100) | 7 (100) |
|  | HN-PGL | 13 (100) | – | 0 (0) | – | 13 (100) | – |
|  | PCC | 12 (11) | 0 (0) | 103 (89) | 4 (100) | 115 (100) | 4 (100) |
| SYP | sPGL | 2 (33) | 1 (14) | 4 (67) | 6 (86) | 6 (100) | 7 (100) |
|  | HN-PGL | 0 (0) | – | 13 (100) | – | 13 (100) | – |
|  | PCC | 7 (6) | 0 (0) | 108 (94) | 4 (100) | 115 (100) | 4 (100) |
| VIM | sPGL | 1 (17) | 3 (46) | 5 (83) | 4 (54) | 6 (100) | 7 (100) |
|  | HN-PGL | 0 (0) | – | 13 (100) | – | 13 (100) | – |
|  | PCC | 26 (23) | 1 (25) | 89 (77) | 3 (75) | 115 (100) | 4 (100) |
| Ki-67 | sPGL | 0 (0) | 0 (0) | 7 (100) | 7 (100) | 7 (100) | 7 (100) |
|  | HN-PGL | 0 (0) | – | 13 (100) | – | 13 (100) | – |
|  | PCC | 15 (13) | 0 (0) | 103 (87) | 4 (100) | 118 (100) | 4 (100) |
| HIF-1α | sPGL | 6 (100) | 6 (100) | 0 (0) | 0 (0) | 6 (100) | 6 (100) |
|  | HN-PGL | 13 (100) | – | 0 (0) | – | 13 (100) | – |
|  | PCC | 109 (97) | 3 (75) | 3 (3) | 1 (25) | 112 (100) | 4 (100) |
| HIF-2α-N | sPGL | 7 (100) | 5 (83) | 0 (0) | 1 (17) | 7 (100) | 6 (100) |
|  | HN-PGL | 8 (67) | – | 4 (33) | – | 12 (100) | – |
|  | PCC | 112 (99) | 4 (100) | 2 (1) | 0 (0) | 114 (100) | 4 (100) |
| HIF-2α-C | sPGL | 4 (57) | 1 (17) | 3 (43) | 5 (83) | 7 (100) | 6 (100) |
|  | HN-PGL | 1 (8) | – | 11 (92) | – | 12 (100) | – |
|  | PCC | 76 (67) | 3 (75) | 38 (33) | 1 (25) | 114 (100) | 4 (100) |
| DEC1/BHLBE40 | sPGL | 2 (33) | 3 (42) | 4 (67) | 4 (58) | 6 (100) | 7 (100) |
|  | HN-PGL | 1 (8) | – | 12 (92) | – | 13 (100) | – |
|  | PCC | 57 (50) | 2 (50) | 58 (50) | 2 (50) | 115 (100) | 4 (100) |

**Table S3.**
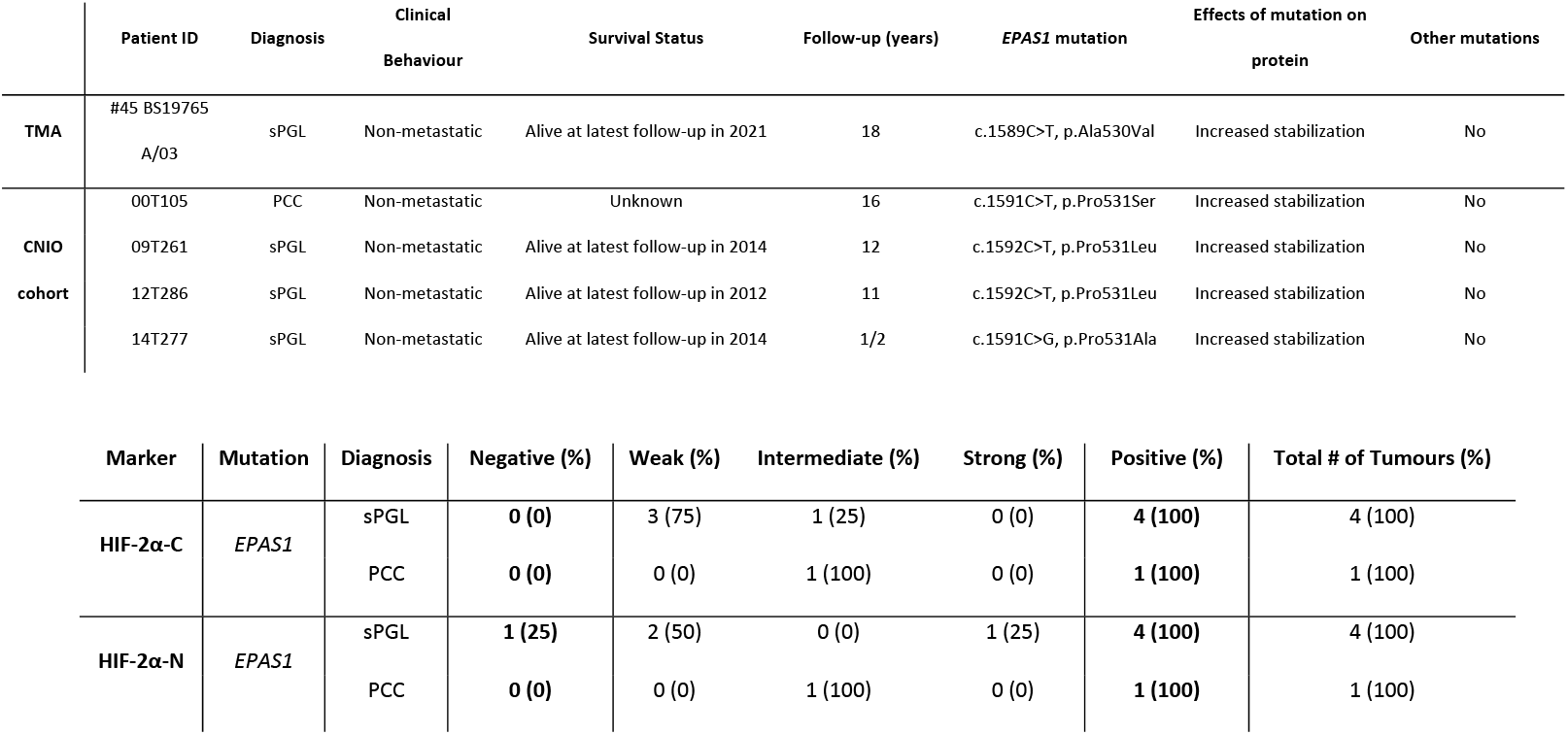
General and staining characteristics of the tumor samples with EPAS1 mutation.

**Table S4.** Causes of death for PPGL patients. Number and fraction (%) of patients dead from metastatic PPGL as compared to all causes of death. PCC, pheochromocytoma; sPGL, sympathetic paraganglioma; HN-PGL, head- and neck paraganglioma. PGL, paraganglioma; PPGL, pheochromocytoma and paraganglioma.

| 10-year survival |  |  |  |  |  |
| --- | --- | --- | --- | --- | --- |
| Cause of Death | Status | sPGL (%) | HN-PGL (%) | PCC (%) | Total (%) |
| All | Alive | 7 (50) | 13 (100) | 104 (85) | 124 (83) |
|  | Dead | 7 (50) | 0 (0) | 18 (15) | 25 (17) |
|  | Total | 14 (100) | 13 (100) | 122 (100) | 149 (100) |
| Metastatic PPGL | Alive | 7 (70) | 13 (100) | 104 (98) | 124 (88) |
|  | Dead | 3 (30) | 0 (0) | 2 (2) | 5 (12) |
|  | Total | 10 (100) | 13 (100) | 106 (100) | 129 (100) |

